# A reverse vaccinology approach identifies putative vaccination targets in the zoonotic nematode *Ascaris*

**DOI:** 10.1101/2022.04.27.489657

**Authors:** Francisco Miguel Dias Evangelista, Arnoud H. M. van Vliet, Scott P. Lawton, Martha Betson

## Abstract

Ascariasis is the most prevalent helminthic disease affecting both humans and pigs and is caused by the roundworms *Ascaris lumbricoides* and *Ascaris suum*. While preventive chemotherapy continues to be the most common control method, recent reports of anthelminthic resistance highlight the need for development of a vaccine against ascariasis. The aim of this study was to use a reverse vaccinology approach to identify potential vaccine candidates for *Ascaris*. Three *Ascaris* proteomes predicted from whole-genome sequences were analysed. Candidate proteins were identified using open-access bioinformatic tools (e.g. Vacceed, VaxiJen, Bepipred 2.0) which test for different characteristics such as sub-cellular location, T-cell and B-cell molecular binding, antigenicity, allergenicity and phylogenetic relationship with other nematode proteins. From over 100,000 protein sequences analysed, four transmembrane proteins were predicted to be non-allergen antigens and potential vaccine candidates. The four proteins are a Piezo protein, two voltage-dependent calcium channels and a protocadherin-like protein, are all expressed in either the muscle or ovaries of both *Ascaris* species, and all contained high affinity epitopes for T-cells and B-cells. The use of a reverse vaccinology approach allowed the prediction of four new potential vaccination targets against ascariasis in humans and pigs. These targets can now be further tested in *in vitro* and *in vivo* assays to prove efficacy in both pigs and humans.

## 1. Introduction

The giant roundworm *Ascaris* is the most prevalent soil-transmitted helminth (STH) infection in humans, being responsible for 0.861 million disability-adjusted life years (DALYs) worldwide (Kyu et al., 2018). In parallel, *Ascaris* infections remain an issue on many pig farms worldwide. This parasite is especially important in farms with lower levels of biosecurity, such as organic farms, and/or in lower to medium income countries, where backyard farming is common (Dold and Holland, 2011; Katakam et al., 2016). Being a zoonotic disease, the close contact between humans and pigs in many highly endemic areas increases the risk *Ascaris* transmission among both hosts (Dold and Holland, 2011), (Easton et al., 2020).

Preventive chemotherapy, bolstered by implementation of water, sanitation and hygiene (WASH) protocols, continues to be the mainstay of ascariasis control in humans, as advocated in the World Health Organization (WHO) roadmap for the control of Neglected Tropical Diseases (NTDs) (World Health Organization, 2020). Reduced efficacy and resistance to benzimidazole drugs have already been reported in human *Ascaris* infections (Furtado et al., 2019; Krücken et al., 2017), and this directly affects the implementation of control protocols in endemic areas, necessitating the development of effective vaccines and vaccination protocols. Vaccination against *Ascaris* has had some degree of efficacy in mouse and pig animal models, but no candidate vaccine has undergone human clinical trials yet (Hotez et al., 2016). The proteins used in the past vaccination assays were either secreted proteins or those retrieved from crude extracts of adult worms. Recently, vaccination with a chimeric protein led to up to 73% larval burden reduction in mice, in contrast to a 99.8% reduction when mice were repeatedly infected with *Ascaris* sterile eggs (de Castro et al., 2021; Gazzinelli-Guimarães et al., 2018; Tsuji et al., 2001, 2004). These results highlight that there are other antigens that could be tested. These antigens could then be incorporated in a multi-component vaccine along with the already known vaccination targets to stimulate a more complete immune response.

Several annotated genomes are now available for *Ascaris lumbricoides* and *Ascaris suum*, which enable the search for putative vaccine candidates using a reverse vaccinology analysis. A reverse vaccinology approach combines genome information and bioinformatic tools for identification of vaccine candidates, and has been successfully applied to other nematodes before, such as *Toxocara canis* and *Trichuris muris* (Salazar Garcés et al., 2020; Zawawi et al., 2020). Such methodology allows researchers to uncover vaccine targets from predicted proteomes by assessing if proteins have useful characteristics. Different analyses, such as protein sub-cellular location and the prediction of B-cell and T-cell epitopes, are examples of steps used in these approaches that help select proteins for further testing. This is especially important when funding is limited, and vaccination trials need to be focused. The aim of this study was to apply an *in silico* methodology to analyse protein sequences predicted from three *Ascaris* annotated proteomes to identify potential new vaccination targets that could be used in vaccination assays against *Ascaris*.

## 2. Materials and methods

### 2.1. Data Retrieval

The annotated proteomes for three assembled genomes of *Ascaris* spp. were retrieved from the WormBase ParaSite database (Howe et al., 2017). The *A. lumbricoides* proteome from BioProject PRJEB4950 (International Helminth Genomes Consortium, 2019) has 23,604 protein sequences. The *A. suum* proteomes from BioProject PRJNA80881 (Jex et al., 2011) and BioProject PRJNA62057 (Wang et al., 2017, 2012) have 18,542 and 57,968 protein sequences, respectively.

### 2.2. Protein subcellular location analysis

The retrieved proteomes were first visualised and analysed with BioEdit v7.2.5 (Hall, 1999). A total of 2,604 protein sequences that included unknown amino acids (aa) (indicated with the symbol ‘X’) were excluded from further analysis as they tend to be a consequence of poor annotations and most of the bioinformatics tools used do not analyse protein sequences with unknown amino acids.

The framework Vacceed v2.1 (Goodswen et al., 2014) was used to identify potential vaccine candidates. The tools employed in Vacceed analysis were: WoLF PSORT v0.2 (Horton et al., 2007), DeepLoc v1.0 (Almagro Armenteros et al., 2017), SignalP v5.0 (José Juan Almagro Armenteros et al., 2019), TargetP v2.0 (Jose Juan Almagro Armenteros et al., 2019), TMHMM v2.0 (Krogh et al., 2001) and Phobius v1.01 (Käll et al., 2004). Proteins were ranked with scores between 0 and 1, from low immunogenicity (final score = 0) to high immunogenicity potential (final score = 1). Higher scores were given to proteins which were predicted to be secreted, signal or transmembrane peptides. The proteins sequences with a Vacceed score of ≥ 0.750 were retrieved (Palmieri et al., 2017) and later tested for epitope binding to CD4+ T helper (Th) cells.

As Vacceed makes use of different bioinformatic tools, there was the need to check how much each tool influences the final protein scores. After retrieving the Vacceed scores from runs including all programs, the process was repeated six times excluding one program per run (i.e. first run with all the bioinformatic tools except TMHMM, second with all the bioinformatic tools except DeepLoc, and so on) (Palmieri et al., 2017). The proteins scores were then compared between the different assessments through a Pearson correlation test using R v3.6.3 (R Core Team, 2020), RStudio v1.2.5033 (RStudio Team, 2019) (with the package *corrplot* (Wei and Simko, 2017)) and IBM SPSS Statistics v26.

### 2.3. CD4+ Th cell binding predictions

The protein sequences retrieved with Vacceed were submitted to the standalone version of the Immune Epitope Database (IEDB) Major Histocompatibility Complex class II (MHC-II) binding predictor v2.22.3 (MHCII-IEDB, available at http://tools.iedb.org/mhcii/). This tool employs neural networks trained on IEDB experimentally validated epitopes to predict and quantify the binding affinity between a given peptide/antigen epitope and a selected MHC-II molecule recognized by CD4+ Th cells. The IEDB-recommended 2.22 prediction method was used, comprising the Consensus approach, NN-align, SMM-align, CombLib and Sturniolo methods, or NetMHCIIpan, if any of the previous methods was not available for the selected MHC-II allele (Fleri et al., 2017). This method was used against the 27 human leukocyte antigen (HLA) allele reference set, covering around 99% of the worlds human population (Greenbaum et al., 2011). The default peptide epitope length of 15 aa with a core of 9 aa was selected. The protein sequences which had epitopes ranked from zero to one (from the maximum of 100) and were simultaneously predicted to bind to all the 27 alleles in the used reference set were selected for further testing. These MHC-II binding predictions were only made for human alleles due to the lack of *in silico* tools that make binding predictions for swine MHC-II alleles.

### 2.4. Allergenicity, antigenicity and function predictions

The protein sequences retrieved with MHCII-IEDB were tested for antigenicity and allergenicity. Protein antigenicity was analysed using IEDB Kolaskar and Tongaonkar antigenicity scale and VaxiJen 2.0 (Doytchinova and Flower, 2007; Kolaskar and Tongaonkar, 1990). The IEDB Kolaskar and Tongaonkar antigenicity scale method, available at http://tools.iedb.org/bcell/, was used with default setting and threshold of 1.000 and the VaxiJen 2.0 tool, server accessible at http://www.ddg-pharmfac.net/vaxijen/VaxiJen/VaxiJen.html, was used with the default threshold of 0.5. Allergenicity was evaluated using AllerTop 2.0 and AllergenFP (Dimitrov et al., 2014, 2013). The AllerTop 2.0 server, https://www.ddg-pharmfac.net/AllerTOP/index.html, and the AllergenFP tool, http://www.ddg-pharmfac.net/AllergenFP/index.html, analyse protein sequences and compare them to a training set of 2,427 known allergens and 2,427 non-allergens using two different algorithms for the better recognition of allergens and non-allergens, respectively for each tool. Proteins that were classified as both potential antigens and non-allergens were considered as good vaccination targets.

The function of the selected proteins, their protein family and tissue location in both *A. lumbricoides* and *A. suum* were analysed based on the most recent genomic and transcriptomic studies (Easton et al., 2020; Wang et al., 2017). These assays were able to map the location where all the genes and, therefore, proteins were transcribed in both *Ascaris* species as well as in what stages of the parasite lifecycle. This information is important as it allows us to infer what roles these proteins could play in the *Ascaris* lifecycle and how useful they would be as components in a vaccine.

### 2.5. CD4+ Th cell binding epitope selection

To further streamline the epitope selection process, we integrated the information gathered using the Phobius and MHCII-IEDB tools to select extracellular CD4+ Th binding epitopes. We used Phobius to assess the extracellular domains of the predicted proteins and identify if the previously predicted epitopes were present in those areas. Proteins that lacked the full 15 aa epitopes in these areas were disregarded. For each protein, the two non-redundant epitopes (without overlapping aa) predicted to bind to the largest combined number of unique alleles in MHCII-IEDB tool analysis were retrieved. These epitopes were submitted to a BLASTp (Basic Local Alignment Search Tool protein) search (available at https://blast.ncbi.nlm.nih.gov/Blast.cgi) to check for identity in humans and pigs. Epitopes with 100% identity to human or pig epitopes were discarded. The BLASTp analysis was conducted with a threshold of e-value=0.05. Each epitope was also submitted to Allertop 2.0 (Dimitrov et al., 2014) to confirm they were not potential allergens. The best expressed protein transcript for each vaccine target, as found in the most recent genomic studies (Easton et al., 2020; Wang et al., 2017), was submitted to Protter to draw the two-dimensional (2D) protein structure using the information retrieved with Phobius (Käll et al., 2004; Omasits et al., 2014).

### 2.6. B-cell linear epitope prediction and selection

The presence of B-cell linear epitopes was assessed using Bepipred 2.0 webserver (available at http://www.cbs.dtu.dk/services/BepiPred/) (Jespersen et al., 2017). Using the same methodology as in the previous step, proteins that had epitopes found in extracellular domains, exposed to the host immunological system, predicted to be non-allergens, and present in most of the protein transcripts of the different genes were considered good vaccination targets. The two highest scored non-allergen epitopes, with a size between 8 and 40 aa, were retrieved for each protein.

### 2.7. Protein phylogenetic analysis

To assess the relationship between the predicted vaccine targets and their orthologues in other nematodes, a phylogenetic analysis of the predicted vaccine targets was performed including predicted orthologues/similar proteins present in annotated genomes of other parasitic nematodes and *Caenorhabditis elegans* available in WormBase ParaSite database (Howe et al., 2017). Proteins were selected according to the list of orthologues provided by WormBase and, if no orthologue was present for a given nematode, a BlastP analysis was carried out and proteins were retrieved according to the combination of query coverage, identity percentage and e-value. The protein sequences were aligned using MUSCLE (http://www.ebi.ac.uk/Tools/muscle/index.html), the phylogenetic analysis was performed with Maximum-likelihood method using the JTT+G substitution model in MEGA-X (Kumar et al., 2018) and the predicted trees were visualized in iTOL (Letunic and Bork, 2021). Nodal support was tested using bootstrap values, which were calculated with 500 replicates.

## 3. Results

### 3.1. Initial protein selection

From the 100,114 protein sequences extracted from the combined *A. suum* and *A. lumbricoides* proteomes, 97,510 were analysed with the Vacceed framework for protein subcellular location. A total of 28.085 proteins were selected for further analysis (Fig.1), from which 5,984 protein sequences were from *A. lumbricoides*, and 5,004 and 17,097 were *A. suum* protein sequences from the two *A. suum* proteomes, BioProject PRJNA80881 and BioProject PRJNA62057, respectively. The BioProject PRJNA62057 has a higher number of proteins in its proteome due to the way it was assembled, where all the expressed transcripts were kept instead of being merged into a single protein sequence. The Vacceed final scores for each protein in the three proteomes are provided in Supplementary Table 1.

**Figure 1.**
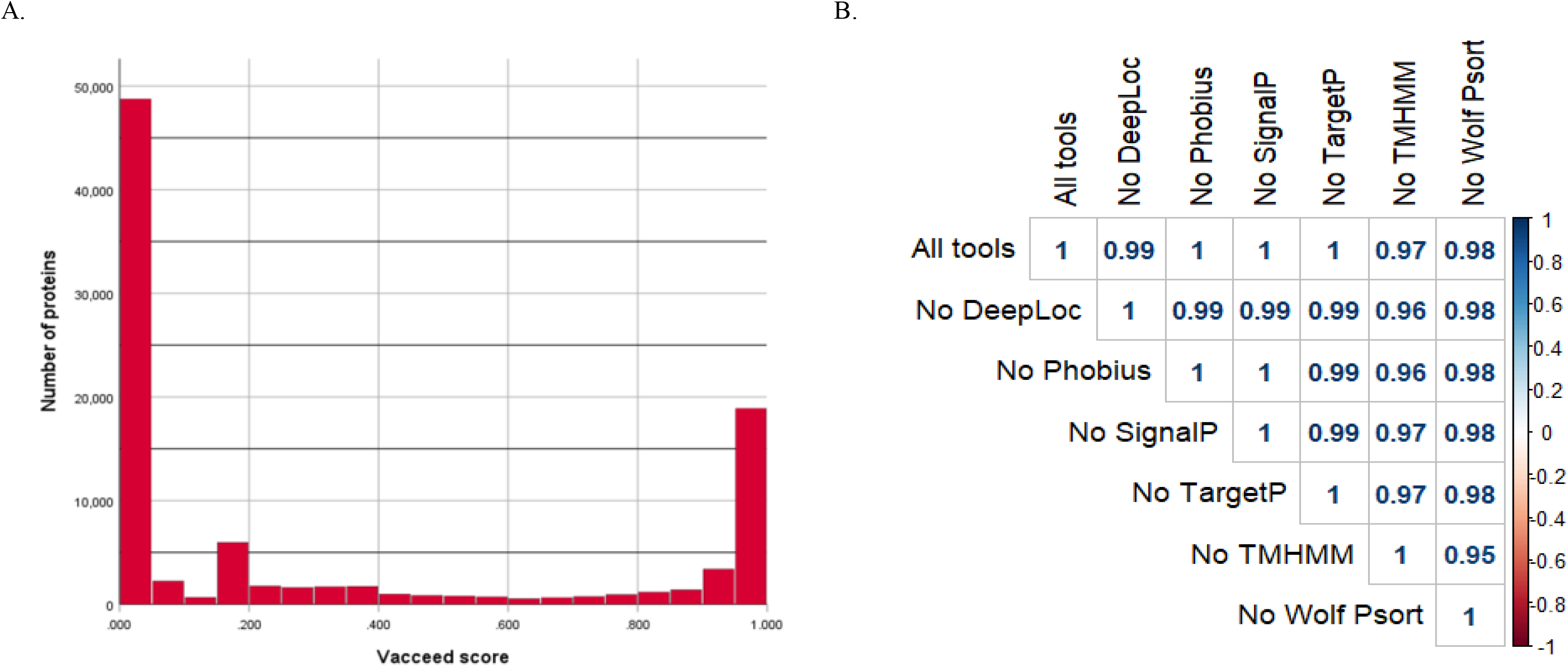
Vacceed assessment. A. Vacceed final scores and respective number of proteins for all analysed protein sequences. B. Pearson correlation plot of the different Vacceed assessments. Each value corresponds to the Pearson correlation coefficient between all the protein sequence scores in two different runs. (All tools = Vacceed with all the used tools: No DeepLoc = Vacceed without the tool DeepLoc; the other labels follow the same principle).

The correlation assessment of the Vacceed framework demonstrated a high correlation between the results of the different Vacceed runs (Figure 1). The removal of TMHMM from the framework resulted in a slightly lowered correlation score when compared with the original Vacceed run (*r*=0.97).

### 3.2. Selection of vaccine targets and identification of epitopes

Before the CD4+ Th cell binding prediction was carried out, a further 2,906 duplicate sequences were identified and removed from the *A. suum* PRJNA62057 dataset. A total of 25,179 protein sequences were then analysed for epitope binding to MHC-II alleles. When analysing the final output, 32 protein sequences had epitopes predicted to bind to all the 27 MHC-II alleles in the reference set with a rank between 0.01 and 1. From this set of 32 protein sequences, 9 proteins belonged to the *A. lumbricoides* proteome, while the other 23 proteins were from the two *A. suum* proteomes, 3 from BioProject PRJNA80881 and 20 BioProject PRJNA62057. Only three of these protein sequences were predicted to be secreted proteins. A table detailing the WormBase protein identifiers and protein aa length for these 32 candidate proteins is provided in Supplementary Table 2.

The analysis with AllerTop 2.0 and AllergenFP did not predict any allergens in the 32 protein sequences. However, antigen prediction tools classified four proteins sequences as non-antigenic, and these proteins were disregarded. The remaining protein sequences that had similar results between their respective orthologues in *A. lumbricoides* and *A. suum* were grouped into five clusters, represented by distinct genes in all three *Ascaris* genomes. The five genes and their respective transcripts are predicted to be responsible for the transcription of four membrane transporters and one cell adhesion protein.

Four genes and respective proteins were selected as promising vaccination targets based on epitope exposure to the host immunological system: “Voltage-dependent T-type calcium channel subunit alpha” (ATtype [WormBase gene identifiers: *GS_24322, ALUE_0000418301* and *AgB13X_g094*]), “Piezo-type mechanosensitive ion channel component” (APiezo [WormBase gene identifiers: *GS_03113, ALUE_0000666901* and *AgR007_g063*]). “Voltage-dependent L-type calcium channel subunit alpha” (ALtype [WormBase gene identifiers: *GS_04697, ALUE_0001482301* and *AgR007_g282*]), and “Protocadherin-like” (AProto [WormBase gene identifiers: *GS_06422, ALUE_0000418601* and *AgB13X_g096*]). According to the most recent transcriptomic data, both ATtype and AProto are highly expressed in the ovaries while APiezo and ALtype are highly expressed in the muscle of adults (Easton et al., 2020; Wang et al., 2017). These proteins were predicted to have both CD4+ Th cell and B cell binding epitopes in extracellular areas (Fig. 2). For each protein, two non-allergen CD4+ Th cell binding epitopes were selected based on the results of the Allertop 2.0, MHCII-IEDB tool and their presence in the extracellular areas of the protein (Table 1). Two B-cell epitopes were also selected for each target. Supplementary Table 3 lists the epitopes found for each predicted vaccine target using MHCII-IEDB that scored between 0 and 1 and the MHC-II alleles they were predicted to bind to (only epitopes that were predicted to bind to two or more MHC-II alleles were retrieved). A workflow diagram of the analysis can be seen in Figure 3.

**Figure 2.**
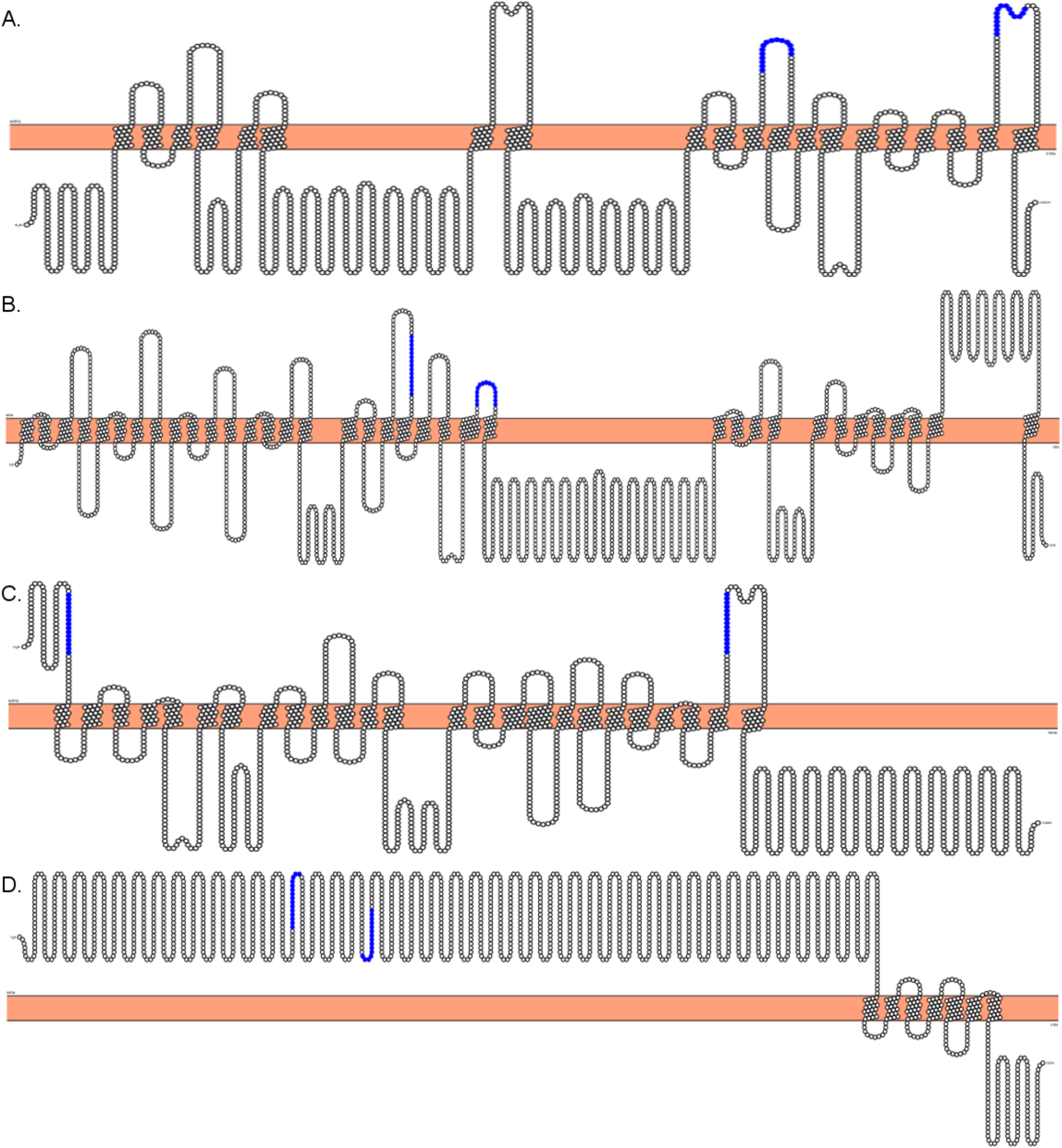
2D representation of the topology of predicted vaccination targets using Phobius predictions. The horizontal bar in the middle represents the membrane, above it is the extracellular domain, and below is the intracellular domain. Highlighted are the Cd4+ Th epitopes chosen to be incorporated in future vaccination assays. A. *“AgB13X_g094_t05”* as the best expressed protein transcript for ATtype. B. *“AgR007_g063_t01”* as the best expressed transcript for APiezo. C. *“AgR007_g282_t05”* as the best expressed protein transcript for ALtype. D. *“AgB13X_g096_t06”* as the best expressed protein transcript for AProto.

**Figure 3.**
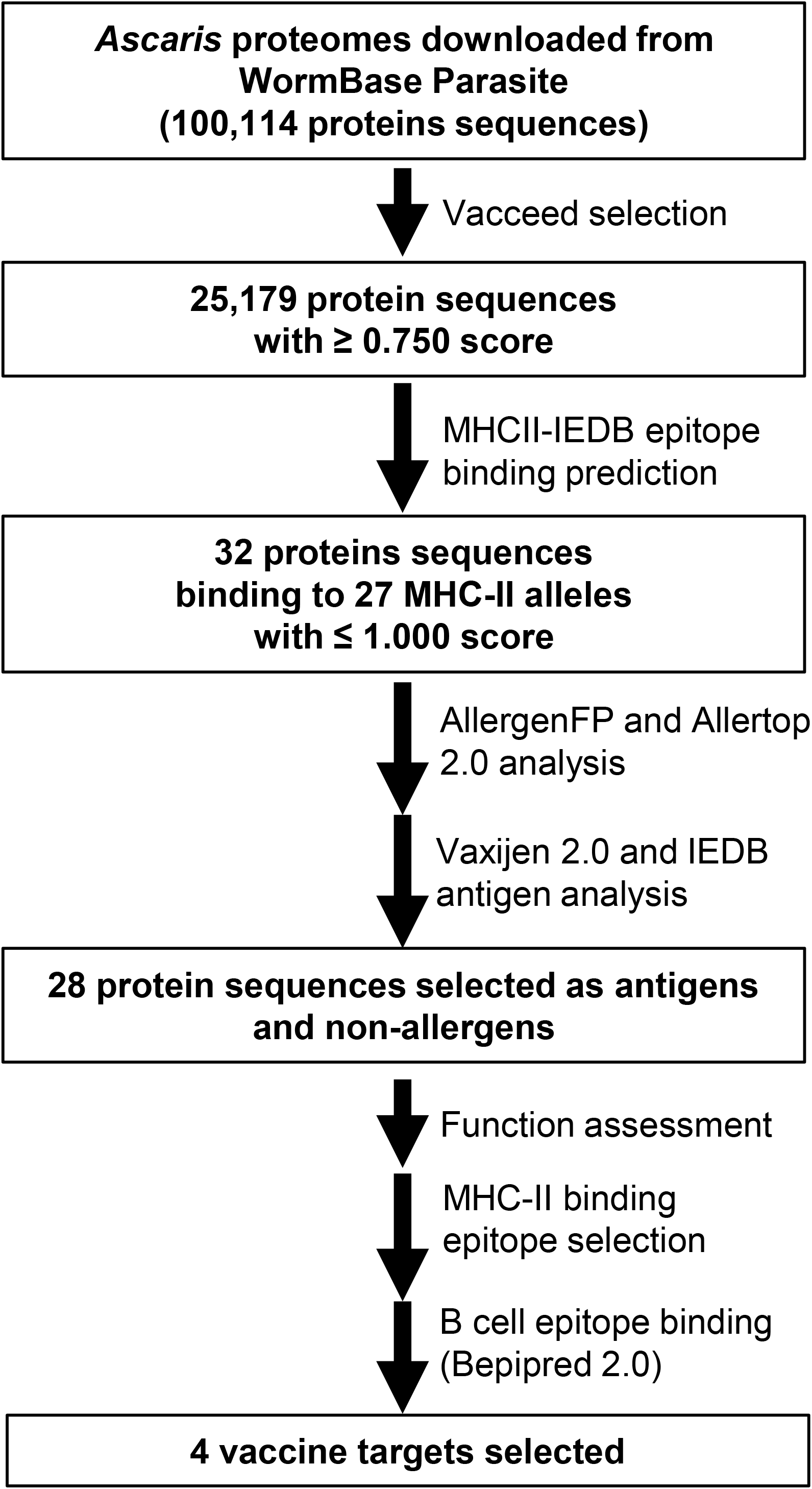
Workflow and summary diagram of the reverse vaccinology approach used in this study to identify and select potential vaccine targets in *Ascaris lumbricoides* and *A. suum*.

**Table 1.**
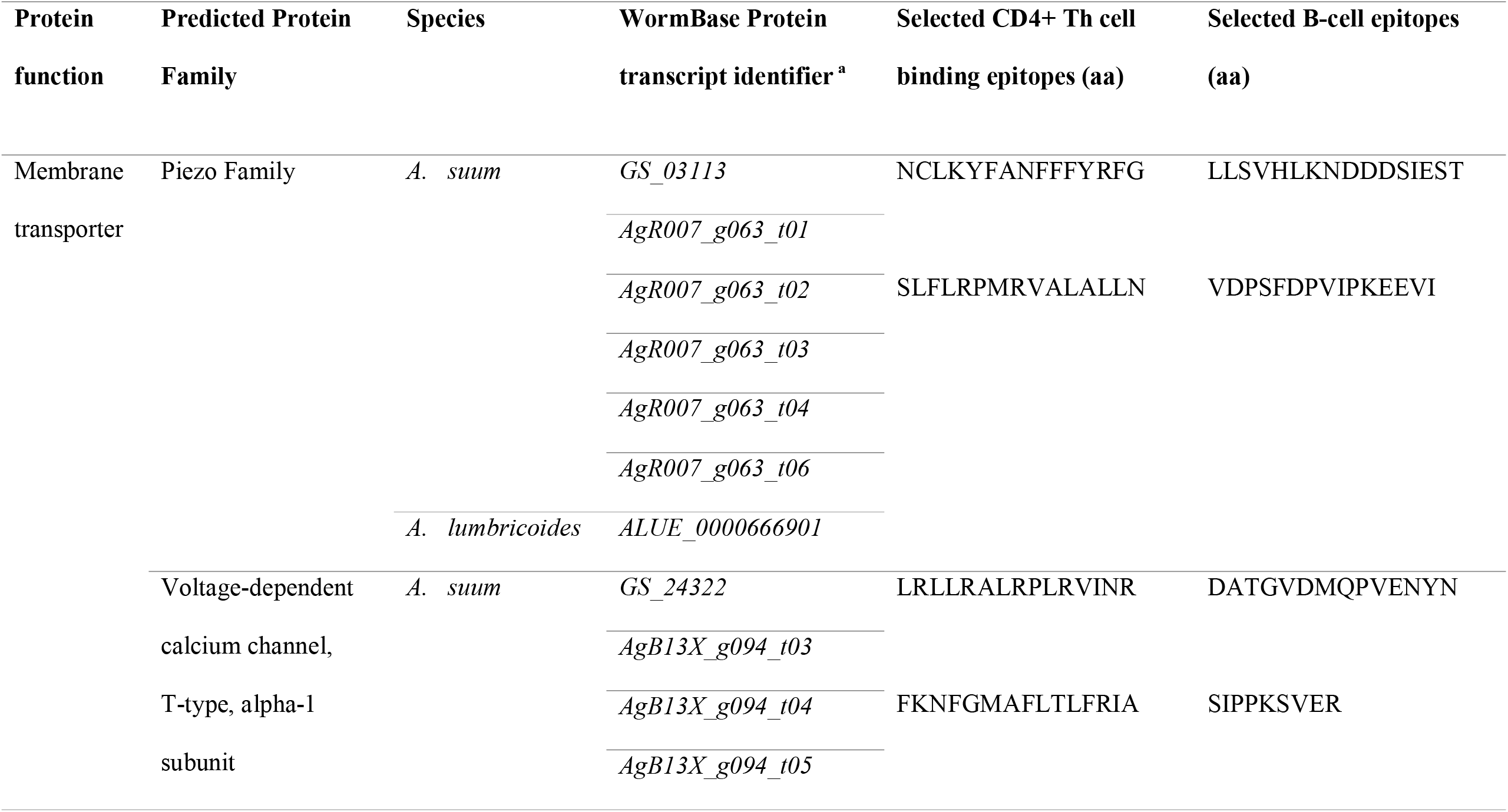
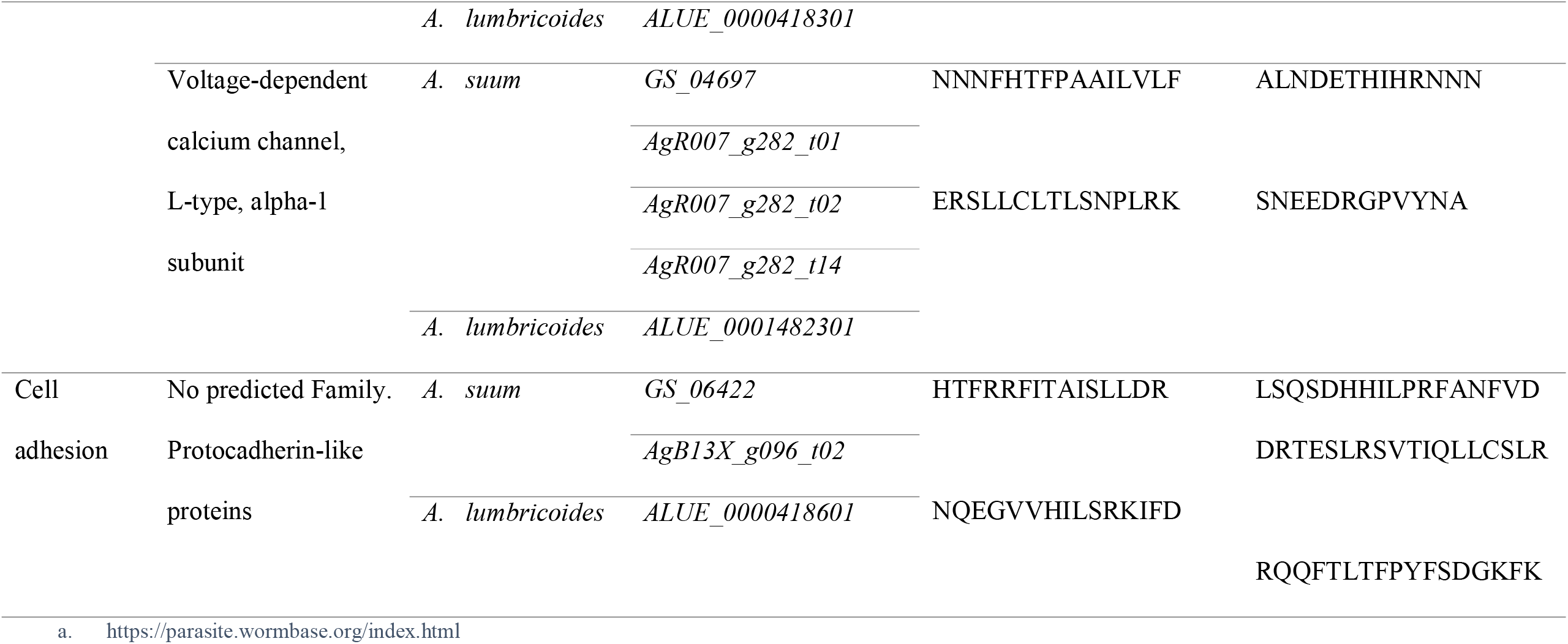
Proteins and respective epitopes identified as potential vaccination targets against *Ascaris lumbricoides* and *Ascaris suum*. Amino acid, aa; *Ascaris suum*, *A. suum*; *Ascaris lumbricoides*, *A lumbricoides*.

### 3.3. Phylogenetic relationships of vaccine targets identified across nematode species

To analyse the potential relationship between the predicted vaccine targets and proteins in other nematodes, orthologs of the selected vaccine targets were retrieved from other nematode proteomes present in WormBase Parasite (Supplementary Table 4), aligned and used for the generation of a phylogenetic tree. There is a general clustering of the proteins within each nematode clade. The exception occurred in the APiezo orthologues where clade III nematodes were separated in two different groups, with the Ascaridomorpha nematodes (e.g. *Ascaris, Toxocara* and *Parascaris*) being more closely related to the nematodes in Clade IV and the Spiruromorpha nematodes (e.g. *Onchocerca, Dirofilaria, Loa, Brugia* and *Wuchereria*) being closer to the Clade V nematodes. These relationships can be seen in Figure 4.

**Figure 4.**
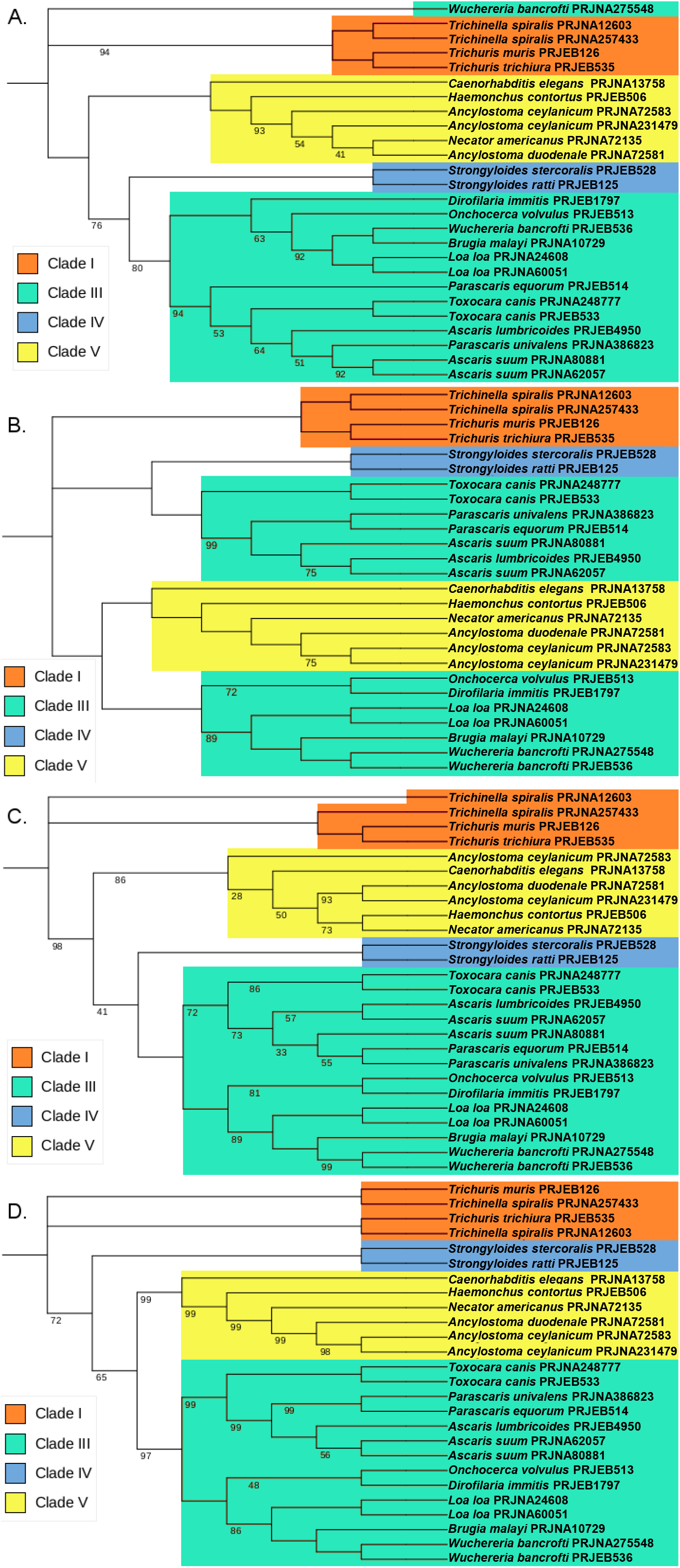
Maximum likelihood phylogenies using the predicted vaccine targets and orthologues found in other nematodes. The trees were inferred using bootstrap values with 500 replicates. Values on the nodes represent the percentage of bootstrap support values and nodes without bootstrap values were supported by 100% of the 500 replicates. Each sequence is identified by the nematode species followed by the respective BioProject (proteins transcripts used in these analyses are individualized in Supplementary Table 4). A. Phylogenetic tree of ATtype orthologues. B. Phylogenetic tree of APiezo orthologues. C. Phylogenetic tree of ALtype orthologues. D. Phylogenetic tree of AProto orthologues.

## 4. Discussion

In this study we have used bioinformatic approaches to identify four different *Ascaris* proteins that could be included in multi-epitope vaccines against human and pig *Ascaris* infections. These genes are highly expressed in two distinct regions of the parasite: the ovaries and early egg stages, in the case of ATtype and Aproto, and in the muscle, the case for APiezo and ALtype (Easton et al., 2020; Wang et al., 2017). While both ATtype and ALtype are predicted to be calcium channels that promote calcium import with muscle and smooth muscle contraction through a voltage mechanism, APiezo is a mechanosensitive calcium channel that in *C. elegans* was shown to affect reproductive tissue development and malfunction (Bai et al., 2020). One interesting relationship shown in Figure 4.B is how APiezo appears to be more closely related between blood feeding parasites, such as *Necator americanus* and filarial parasites such as *Loa loa*, suggesting a role in adaptation to contact with blood. The AProto protein is the most unique as to our knowledge, as protocadherins have not been reported before in nematodes. The structural domains appear to be more closely related to that of Flamingo/Stan cadherins due to the presence of both laminin-G and EGF-like receptors with seven transmembrane domains (with these last highlighted in Figure 2D) (Easton et al., 2020; Hardin et al., 2013). Being highly transcribed in the ovaries and early egg stages, while predicted to be responsible for homophilic cell adhesion, AProto could have a role in early oocyte development. Although present in regions usually considered difficult recognize by the host immune system, such as the parasite’s muscle and ovaries, an IgG immunoblot assay showed that this is possible, as proteins highly transcribed in the muscle, ovaries and intestines of the parasite were recognized by serum from pigs infected with *Ascaris* (González-Miguel et al., 2014). It is also interesting to highlight the close relationship between the predicted vaccine targets and the orthologues in other nematodes, suggesting that these orthologues might also be useful in the control of the respective species.

Previous attempts to generate a vaccine against ascariasis used crude extracts, recombinant proteins and, more recently, chimeric proteins (de Castro et al., 2021; Gazzinelli-Guimarães et al., 2018; Tsuji et al., 2004, 2001). The highest lung larvae burden reduction achieved using a multi-epitope or recombinant protein vaccine was 73.5%, when using a chimeric protein containing B-cell epitopes of the As14, As16 and As37 *Ascaris* proteins (de Castro et al., 2021). In comparison, vaccination against trichuriasis in mice resulted in up to 97% reduction of adult nematodes (Gomez-Samblas et al., 2017). The vaccine against *Trichuris* is based on recombinant proteins and showed efficacies vastly superior to similar vaccines against ascariasis. With vaccine development against ascariasis lagging behind other parasitic diseases, there is a need to discover other antigens to be tested as vaccine candidates. The reverse vaccinology approach used in our study is based on genomic and proteomic data to predict which proteins may be usable as vaccine candidates prior to new *in vivo* studies. This methodology allows researchers to focus down vaccination assays to a smaller set of proteins, effectively reducing costs and time. This reverse vaccinology approach has been successfully applied to identify vaccination targets for other nematodes, for example *T. canis* (Salazar Garcés et al., 2020) and *T. muris* (Zawawi et al., 2020).

Our reverse vaccinology analysis used all the proteins predicted in the three *Ascaris* proteomes. Previous *in silico* studies on vaccine candidate prediction in *Ascaris* focused exclusively on secreted proteins (Ebner et al., 2020). The workflow applied in this study allows the testing of all the proteins predicted from a genome analysis, without automatically excluding non-secreted proteins. In a recent study, an *A. lumbricoides* multi-epitope vaccine candidate was developed using *in silico* methodology and proteins were selected based on their binding to the HLA-DRB1*07:01 and HLA-DRB1*15:01 MHC-II alleles (Kaur et al., 2021). Although these MHC-II alleles are known to recognise *Ascaris* antigens in humans, they only cover up to 15% of the human population in areas where human ascariasis is endemic (Ebner et al., 2020). This might prove detrimental in *in vivo* studies that cover population that do not have these MHC-II alleles, limiting its usefulness.

Each of the three genomes included in the analysis were predicted to have over 5,000 proteins that could be further investigated as good vaccine targets according to Vacceed, which corresponds to 25-29% of all the proteins present. As the number of secreted proteins in *Ascaris* is predicted to be 254 proteins (Ebner et al., 2020), this suggests that the number of potential vaccination targets might be vastly superior to the ones that are usually investigated in these species and other helminths. The final candidates are all predicted to be non-secreted proteins. This contrasts with most of the previously studied vaccine candidates, except for the muscle membrane-bound As37 protein (Versteeg et al., 2020). As37 was not predicted to be a good target in Vacceed (with scores of 0.001 in all three genomes), showing some limitations to the method we used. However, our predictions and the protection achieved with the As37 recombinant protein support the idea that only targeting secreted proteins can be detrimental to the selection of good vaccination targets in *Ascaris*. Recent work in *Toxocara canis*, a parasite of the same family as *Ascaris*, showed that membrane proteins are capable to induce protection in a mouse model, reinforcing the idea that it is an error to disregard these proteins when looking for vaccination targets in nematodes (Salazar Garcés et al., 2020).

The underlying host’s immune responses against *Ascaris* are still only partially understood (Zawawi and Else, 2020). This is a disadvantage when selecting the right tools to predict which proteins could be useful as vaccination targets. MHC-II molecules appear to have a prominent role in the control of nematode infections in mammals, including *A. lumbricoides* infections in humans (Garamszegi and Nunn, 2011; Zawawi and Else, 2020). This role makes the discovery of epitopes that bind to these molecules a priority for the design of multi-epitope/subunit-based vaccines. The MHCII-IEDB tool was chosen for this purpose as it was used in selecting epitopes for vaccination assays against other nematodes and is one of the most accurate tools available (Ebner et al., 2020; Zawawi et al., 2020). A reference set of 27 different alleles was chosen due to the fact that heterogenicity throughout the human population leads to different immune responses(Ebner et al., 2020). Thus, we wanted to selected proteins that would be able to induce a helpful immune reaction in a large portion of the population, and not just focus on one specific allele. Unfortunately, this biases towards selecting larger proteins as they contain a larger number of epitopes. These predictions were made only for human alleles due to unavailability of similar open-access bioinformatic tools for swine MHC-II alleles. This is a limitation of the reverse vaccinology approach to discovering vaccine targets in pigs. Bepipred 2.0 predicted the presence of B cell binding epitopes in same proteins which would enhance the host immune response against these proteins in their native form in the parasite.

Whilst the proteins we selected have been predicted to be useful and have similarities to other proteins tested in vaccination assays against other nematodes, there is still the need to test them *in vitro* and *in vivo*. The next step should be to confirm *in vitro* that both humans and pigs are able to recognize these proteins targets and their respective epitopes. Should these proteins prove useful in stimulating an immune reaction in the host, we propose that the proteins and respective epitopes identified in this work should be incorporated into a multi-epitope vaccine which, ideally, would include CD4+ Th cell and B-cell binding epitopes from other proteins, such as As14, As16 and As37. Ideally, such a vaccine should be tested using a pig model to assess its potential effect on both larvae and adult *Ascaris*, impossible to assess using a mouse model. This is more relevant when testing for the utility of incorporating ATtype and AProto epitopes due to protein higher expression in the ovaries of adult female *Ascaris* (Easton et al., 2020; Wang et al., 2017). Should these proteins not prove useful, a protocol optimization should be done to identify new targets so that, in the future, it might also be applied to other species that are in need of further studies.

In conclusion, this study highlights the role that reverse vaccinology and *in silico* methodology can play in identification of vaccine candidates for parasitic diseases. The genome-wide approach, without bias towards secreted proteins, led to the prediction of four novel candidates that were not identified in previous studies but show the promise of promoting a useful immune response in vaccination assays against *A. lumbricoides* and *A. suum*. These proteins should now be tested *in vitro* and in combination with already known vaccine targets. Ultimately, the findings of this study will support the future development of a vaccine against both ascariasis in humans and pigs, thus promoting the health of both populations by reducing the need to use Mass-Drug administrations and decreasing the risk of anthelmintic resistance appearing.

## Acknowledgements

The authors would like to thank Dr. Stephen Goodswen for assistance and troubleshooting with the Vacceed tool and Dr. Pritesh Taylor for help with computer resources. Graphical abstract created with BioRender.com.

## Author contributions

Concept and design: FE, AV, SL and MB

Data acquisition and analysis: FE

Data interpretation: FE, AV, SL and MB

Manuscript draft: FE

Critical review and editing of the manuscript: FE, AV, SL and MB

---

Supplementary Table 1 – The Vacceed final scores for each proteome in the three *Ascaris* proteomes.

Supplementary Table 2 - Protein sequences predicted to have epitopes that bind to all 27 MHC-II alleles, used as a reference set, with a score between 0.01 and 1 in the MHCII-IEDB tool.

Supplementary Table 3 - The epitopes found for each predicted vaccine target using MHCII-IEDB that scored between 0 and 1 and the MHC-II alleles they were predicted to bind to (only epitopes that were predicted to bind to two or more MHC-II alleles were retrieved).

Supplementary Table 4 – List of orthologues used in the phylogenetic analysis.

