## Supplemental Table S2 for "A reverse vaccinology approach identifies putative vaccination targets in the zoonotic nematode *Ascaris*"

Supplementary Table 2 - Protein sequences predicted to have epitopes that bind to all 27 MHC-II alleles, used as a reference set, with a score between 0.01 and 1 in the MHCII-IEDB tool.

| Species | Bioproject | WormBase Protein transcript identifier | Protein transcript length (aa) | Phobius sub-cellular location |
| --- | --- | --- | --- | --- |
| A. *lumbricoides* | PRJEB4950 | ALUE_0000401101 | 3420 | Transmembrane |
|  |  | ALUE_0000418301 | 2196 | Transmembrane |
|  |  | ALUE_0000418601 | 2591 | Transmembrane |
|  |  | ALUE_0000827601 | 3725 | Transmembrane |
|  |  | ALUE_0000834301 | 3758 | Transmembrane |
|  |  | ALUE_0000985601 | 2181 | Transmembrane |
|  |  | ALUE_0001005801 | 1141 | Transmembrane |
|  |  | ALUE_0001162401 | 2044 | Transmembrane |
|  |  | ALUE_0001794801 | 774 | Transmembrane |
| A. *suum* | PRJNA80881 | GS_03113 | 2294 | Transmembrane |
|  |  | GS_05892 | 3963 | Transmembrane |
|  |  | GS_21459 | 1022 | Transmembrane |
|  | PRJNA62057 | AgB13X_g094_t03 | 2057 | Transmembrane |
|  |  | AgB13X_g094_t04 | 2060 | Transmembrane |
|  |  | AgB13X_g094_t05 | 1862 | Transmembrane |
|  |  | AgB13X_g096_t02 | 2637 | Transmembrane |
|  |  | AgR002_g353_t01 | 2173 | Transmembrane |
|  |  | AgR007_g063_t01 | 2612 | Transmembrane |
|  |  | AgR007_g063_t02 | 2661 | Transmembrane |
|  |  | AgR007_g063_t03 | 2670 | Transmembrane |
|  |  | AgR007_g063_t04 | 2682 | Transmembrane |
|  |  | AgR007_g063_t06 | 2666 | Transmembrane |
|  |  | AgR007_g282_t01 | 1891 | Transmembrane |
|  |  | AgR007_g282_t02 | 1889 | Transmembrane |
|  |  | AgR007_g282_t14 | 1770 | Transmembrane |
|  |  | AgR028_g099_t06 | 2170 | Secreted |
|  |  | AgR028_g099_t07 | 2631 | Secreted |
|  |  | AgR035X_g027_t05 | 3668 | Transmembrane |
|  |  | AgR035X_g062_t03 | 3452 | Secreted |
|  |  | AgR052_g054_t02 | 1386 | Transmembrane |
|  |  | AgR052_g054_t03 | 1412 | Transmembrane |
|  |  | AgR052_g054_t04 | 1384 | Transmembrane |
